# Genomic insights into adaptations of TMA-utilizing methanogens to diverse habitats including the human gut

**DOI:** 10.1101/2020.09.17.302828

**Authors:** Jacobo de la Cuesta-Zuluaga, Timothy D. Spector, Nicholas D. Youngblut, Ruth E. Ley

## Abstract

Archaea of the order *Methanomassiliicoccales* use methylated-amines such as trimethylamine as a substrate for methane production. They form two large phylogenetic clades and reside in diverse environments, from soil to the human gut. Two genera, one from each clade, inhabit the human gut: *Methanomassiliicoccus*, which has one cultured representative, and “*candidatus* Methanomethylophilus”, which has none. Questions remain regarding their distribution across different biomes and human populations, their association with other taxa in the human gut, and whether host genetics correlate with their abundance. To gain insight into the *Methanomassiliicoccales*, and the human-associated members in particular, we performed a genomic comparison of 72 *Methanomassiliicoccales* genomes and assessed their presence in metagenomes derived from the human gut (n=4472 representing 25 populations), nonhuman animal gut (n=145) and nonhost environments (n=160). Our analyses showed that all taxa are generalists: they were detected in animal gut and environmental samples. We confirmed two large clades, one enriched in the gut, the other enriched in the environment, with notable exceptions. Genomic adaptations to the gut include genome reduction, a set of adhesion factors distinct from that of environmental taxa, and genes involved in the shikimate pathway and bile resistance. Genomic adaptations differed by clade, not habitat preference, indicating convergent evolution between the clades. In the human gut, the relative abundance of *Methanomassiliicoccales* correlated with trimethylamine-producing bacteria and was unrelated to host genotype. Our results shed light on the microbial ecology of this group may help guide *Methanomassiliicoccales-based* strategies for trimethylamine mitigation in cardiovascular disease.

**Importance:** *Methanomassiliicoccales* are a lesser known component of the human gut microbiota. This archaeal order is composed of methane producers that use methylated amines, such as trimethylamine, in methane production. This group has only one cultured representative; how they adapted to inhabit the mammalian gut and how they interact with other microbes is largely unknown. Using bioinformatics methods applied to DNA from a wide range of samples, we profiled the relative abundances of these archaea in environmental and host-associated microbial communities. We observed two groups of *Methanomassiliicoccales*, one largely host-associated and one largely found in environmental samples, with some exceptions. When host-associated, these archaea have a distinct set of genes related to adhesion and possess genes related to bile resistance. We did not detect *Methanomassiliicoccales* in all human populations tested but when present, they are correlated with *Bacteria* known to produce trimethylamine. Since trimethylamine is linked to cardiovascular disease risk, these intriguing Archaea may also be involved.

## Introduction

*Archaea* generally make up a tenth or less of the biomass of the human gut microbiota; however, they are widely prevalent and occupy a unique metabolic niche, utilizing by-products of bacterial metabolism as substrate for methanogenesis (1). The most widespread methanogens in the human gut are members of the order *Methanobacteriales.* These include *Methanobrevibacter smithii*, which uses CO_2_, formate and H_2_ as substrates for methane production (2), and *Methanosphaera stadtmanae*, which consumes methanol and H_2_ (3). Through the process of methane formation, *Archaea* decrease partial pressures of H_2_, thereby potentially increasing the energetic efficiency of primary fermenters and the production of short-chain fatty acids (4). Members of *Methanobacteriales* are the dominant species of the human gut archaeome (1, 5).

A second archaeal lineage, the order *Methanomassiliicoccales*, is also found within the human gut microbiota, yet its members are less well characterized than those of *Methanobacteriales.* Members of the order *Methanomassiliicoccales*, including human-derived *Methanomassiliicoccus luminyensis, “candidatus* Methanomassiliicoccus intestinalis” and *“candidatus* Methanomethylophilus alvus”, perform H_2_-dependent methylotrophic methanogenesis as sole energy source (6–8). Their genomes encode several methyltransferases and associated proteins used to reduce methylamines and methanol to methane. Studies based on 16S rRNA and *mcrA* gene diversity analysis indicate that the order *Methanomassiliicoccales* is made up of two large clades, which mostly group species that have either a free living (FL) or host associated (HA) lifestyle (9, 10). Based on analyses of the genomes from three human-derived species from both clades, Borrel *et al.* (11) suggested each clade colonized the mammalian gut independently. Members of the HA clade, including the human-associated *“ca.* M. alvus*”*, might be expected to show adaptations similar to other methanogens from the gut microbiota (12, 13). How members of the FL clade, including the human-associated *M. luminyensis* and *“ca.* M. intestinalis”, have converged on the gut niche remains to be explored.

A better understanding of the ecology of *Methanomassiliicoccales* may be of interest to human health, as they can utilize mono-, di-, and trimethylamine (TMA) as substrate for methanogenesis in the gut (14). TMA, a by-product of the bacterial metabolism of carnitine, choline, and other and choline-containing compounds, is absorbed by the host and transformed in the liver into trimethylamine N-oxide (TMAO) (15). In turn, circulating TMAO inhibits cholesterol transport and promotes its accumulation in macrophages, inducing the formation of artherosclerotic plaques (16). Decreasing TMA levels in the gut, and reducing circulating TMAO levels, has been proposed as a therapeutic strategy for cardiovascular disease (17). One way to use the gut microbiome to this end would be to boost levels of *Methanomassiliicoccales* (18). To accomplish this goal requires a deeper understanding of its ecology.

Here, we conducted a comparative analysis of 71 *Methanomassiliicoccales* genomes, together with an additional metagenome-assembled genome (MAG) corresponding to a strain of “*ca.* M. alvus*”*, which we retrieved by metagenome assembly of gut samples from subjects of the TwinsUK cohort (19). We used 305 publically available metagenomes to assess the prevalence of taxa across various habitat types. While the two large clades grouping host-associated (HA) and free-living (FL) taxa, are generally enriched in host-associated and environmental metagenomes, a few exceptions stand out. Our results showed that the repertoire of adhesion proteins encoded by the genomes of taxa from each clade differed. Genes involved in bile resistance and the shikimate pathway are likely involved in the adaptation to the gut environment of members of the HA clade, but not for the FL clade. Thus, gut-adapted members converged on life in the gut using different genomic adaptations. *Methanomassiliicoccales* genera present in the human gut positively correlate with TMA-producing bacteria.

## Materials and Methods

### Genome annotation and phylogenomic tree reconstruction

We downloaded 78 available genomes belonging to the order *Methanomassiliicoccales* from the NCBI assembly database (https://www.ncbi.nlm.nih.gov/assembly) as available in June 2018, and used CheckM to assess their quality. For subsequent analyses, we included 71 substantially complete genomes (completeness ≥70 %) with low contamination (contamination <5 %) (20), plus an additional high-quality metagenome-assembled genome (MAG) corresponding to *“candidatus* Methanomethylophilus alvus” (see *supplementary methods* and table S1). Gene calling, proteome prediction and annotation was performed on each genome using Prokka 1.12 (21). Details of each genome, including the original source of isolation, can be found in (table S1).

Using PhyloPhlAn 0.26 (22), we constructed a maximum-likelihood phylogenomic tree using a concatenated alignment of multiple universally distributed single copy marker genes of 72 publicly available genomes from the order *Methanomassiliicoccales.* Of these, one was retrieved from pure culture, 6 were obtained from enrichment cultures and 64 were MAGs. We included an additional MAG retrieved from human gut metagenomes corresponding to “*ca*. M. alvus” (*supplementary results*). Briefly, universal markers were obtained from the translated amino acid sequences of the included genomes, aligned using mafft 7.3 (23) and concatenated into a single sequence. We then used the concatenated alignment to reconstruct an maximum-likelihood phylogenetic tree using RAxML 8.1 (24); branch support was estimated by 1000 bootstrap iterations and the tree was rooted by including members of the order *Thermoplasmatales* as outgroup, namely *Thermoplasma acidophilum* DSM 1728 (GenBank assembly accession: GCA_000195915.1), *Picrophilus oshimae* DSM 9789 (GCA_900176435.1), *Ferroplasma acidarmanus* fer1 (GCA_000152265.2), *Acidiplasma aeolicum* (GCA_001402945.1) and *Cuniculiplasma divulgatum* (GCA_900090055.1). We used iTOL (25) to visualize the tree.

### Abundance of Methanomassiliicoccales in environmental and animal gastrointestinal metagenomes

We retrieved 305 metagenome samples of gastrointestinal and environmental origin (26) sequenced using the Illumina HiSeq platform (table S2). Sequences were then downloaded from the Sequence Read Archive (SRA) and quality-controlled (see *supplementary methods*). To avoid the issue of multiple mapping, we dereplicated the 72 genomes at a species-level threshold (95 % ANI) using dRep, resulting in 29 representative genomes. Next, we quantified the abundance of dereplicated *Methanomassiliicoccales* genomes in these samples using KrakenUniq v.0.5.8 (27). Statistical analyses were performed using R v.3.5.1 (28). We estimated the enrichment of each representative *Methanomassiliicoccales* on host or environmental metagenomes using DESeq2 (29) on sequence counts and classifying metagenome samples as either host-derived or environmental. We applied hierarchical clustering using Ward’s method on the log-fold-change of environmental vs gastrointestinal enrichment of each taxon and calculated the cophenetic correlation with the phylogenomic tree using the ape package of R (30).

### Comparative genomics

The predicted proteome of each included genome was used to to assign orthology clusters using panX 1.6.0 (31). We used InterProScan (32) and eggNOG mapper 1.0.3 (33) with DIAMOND 0.8.36 (34) against the optimized archaeal database to improve the annotation of gene clusters. Phylogenetic signal of genome characteristics and gene cluster presence was tested using the phylosignal package of R with the local indicator of phylogenetic association (LIPA) (35).

The R package micropan (36) was used to create a principal component analysis (PCA) of gene cluster presence. We compared the gene cluster content between clades to determine gene clusters enriched on clades FL or HA using phylogenetic ANOVA using the R package phytools (37). To reduce the number of comparisons we first removed low frequency gene clusters by filtering those with near zero variance. The above analysis was repeated by comparing gene cluster content between taxa significantly enriched on gut or environmental samples, prior removal of taxa not significantly enriched in either biome class. We adjusted P values with the Benjamini-Hochberg method.

We assessed the presence of eukaryote-like proteins (ELPs) (38) by combining the counts of gene clusters classified by InterProScan as any of the following: Sel1 containing proteins (Sel1), Listeria-Bacteroides repeat containing proteins (List-Bact), tetratricopeptide repeats (TPRs), Ankyrin repeats (ANKs), Leucine-rich repeats (LRRs), Fibronectin type III (fn3) domains, Laminin G domain, Bacterial Ig-like domains, YadA-like domain (Yersinia adhesin A), TadE-like domain or Invasion protein B (ialB). Likewise, we characterized the presence of parallel beta-helix repeat-containing proteins, also known as adhesin-like proteins (ALPs).

### Characterization of Methanomassiliicoccales distribution across human populations

We obtained sample metadata from publicly available studies using the curatedMetagenomicData v.1.17.0 package of Bioconductor (39). Samples were selected according to the following criteria: i) shotgun gut metagenomes sequenced using the Illumina HiSeq platform with a median read length > 95 bp; ii) with available SRA accession; iii) labeled as adults or seniors, or with a reported age ≥ 18 years; iv) without report of antibiotic consumption (i.e. no or NA); v) without report of pregnancy (i.e. no or NA); vi) non-lactating women (i.e. no or NA); vii) without report of gangrene, pneumonia, cellulitis, adenoma, colorectal cancer, arthritis, Behcet’s disease, cirrhosis or inflammatory bowel disease. Only forward reads were downloaded and processed. A total of 4472 samples from 34 independent studies were downloaded from the SRA between December 2019 and February 2020 (table S3) and quality controlled as described in *supplementary methods.*

Reads were classified using Kraken v.2.0 (40) and a Bayesian re-estimation of the species-level abundance of each sample was then performed using Bracken v.2.2 (41). We utilized custom databases created using the Struo pipeline (42) based on GTDB release 86 (available at http://ftp.tue.mpg.de/ebio/projects/struo/). Taxa with <100 reads in a given sample were considered as absent. We obtained complete taxonomic annotations from NCBI taxIDs with TaxonKit 0.2.4 (https://bioinf.shenwei.me/taxonkit/). To determine the cooccurrence patterns of the detected *Methanomassiliicoccales* in the human gut we used the cooccur package of R (43); to determine their coabundance patterns, we calculated the proportionality of taxa abundance (*rho*) with the propr package (44). The lme4 and lmerTest R packages (45) were used to fit linear mixed effects models to test differences of *Methanomassiliicoccales* genera log-transformed abundance by westernization status, age and gender with F-tests and P-values determined via the Satterthwaite’s method (ANOVA Type II sum of squares). Similarly, we employed binomial linear mixed models to test differences of *Methanomassiliicoccales* genera prevalence.

We assessed the heritability of *Methanomassiliicoccales* taxa by comparing relative abundances within 153 monozygotic (MZ) and 200 dizygotic (DZ) twin pairs using the taxonomic profiles of 706 gut metagenome samples from the United Kingdom Adult Twin Registry (TwinsUK) (19, 46, 47) with a sequencing depth >5 million reads/sample. We aggregated abundances at the genus level and removed genera with a prevalence <5 %. Absolute read counts were transformed using the Yeo-Johnson transformation and adjusted by BMI, sex and sequencing depth (19, 46). For each genus, we calculated the intraclass correlation coefficient (ICC) in MZ and DZ twins with the irr package of R, and adjusted P-values for multiple comparisons using the Benjamini-Hochberg method. As control we compared the mean ICC across all taxa between MZ and DZ twins using the Mann-Whitney test, and by assessing the ICC of specific taxa known previously reported as heritable in the same population (*Methanobrevibacter, Faecalibacterium, Christensenella* and *Bifidobacterium*) (46, 48). We carried a sensitivity analysis by repeating these analyses on a subset of 394 samples (80 MZ and 117 DZ twin pairs) with a sequencing depth of >12 million reads/sample.

### Data and code availability

The metagenomic sequence data generated during this study have been deposited in the European Nucleotide Archive with accession ID PRJEB40256. The jupyter notebooks with analysis code are available at https://github.com/leylabmpi/Methanomassilii. The “*candidatus* Methanomethylophilus alvus” MAG here generated can be found at http://ftp.tue.mpg.de/ebio/projects/Mmassilii

## Results

### Genome-based phylogeny confirms two large Methanomassiliioccales clades

Based on whole-genome phylogenetic analysis, the order *Methanomassiliicoccales* forms two clades with robust support (figure 1). This phylogeny is in agreement with previously reported phylogenies based on 16S rRNA and mcrA genes (9, 49, 50). A third distal clade was formed by two closely related MAGs generated in a recent massive metagenome assembly effort (51), which we labeled external (EX; figure 1). We use the terminology of Borrel *et al.*, (52): the clade including *Methanomassiliicoccus* is labeled free-living (FL), and the clade containing “*candidatus* Methanomethylophilus” host-associated (HA).

**Figure 1.**
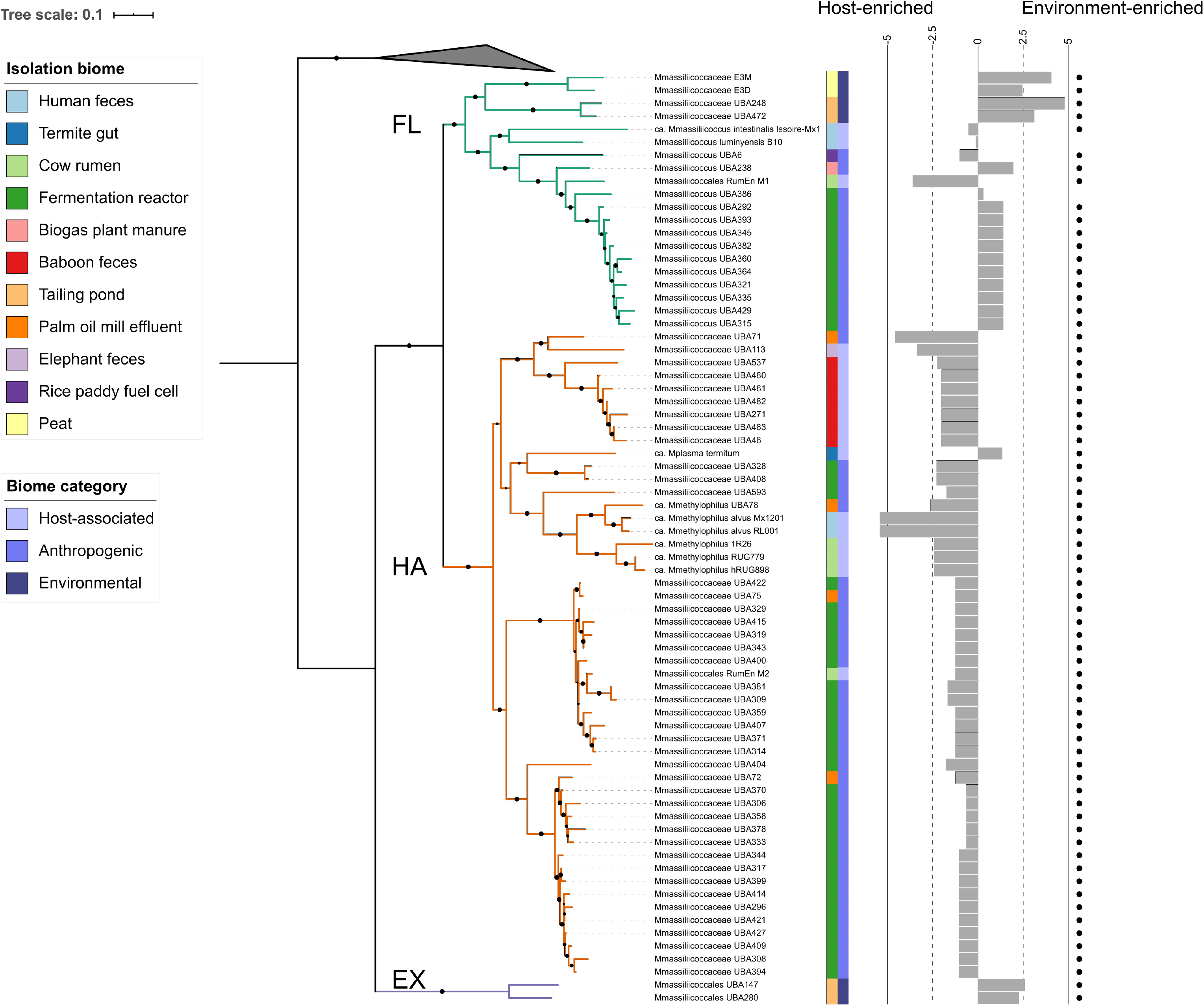
The order *Methanomassiliicoccales* forms two large clades that loosely follow the source of isolation. A maximum likelihood phylogeny of concatenated single-copy marker genes. The gray triangle corresponds to *Thermoplasma acidophilum, Picrophilus oshimae, Ferroplasma acidarmanus, Acidiplasma aeolicum* and *Cuniculiplasma divulgatum*; outgroup taxa from class *Thermoplasmata.* Black circles indicate bootstrap values of > 80 (of 100 bootstrap permutations), and branch color represents the clade. Colored strips show the source of isolation of each of the included genomes and the general category to which the source belongs. Bar plots show the genome abundance enrichment in gut metagenome samples compared to environmental samples calculated using DESeq2; dots indicate taxa with significant enrichment in either host or environmental biome (Adj. P < 0.05). The scale bar represents the number of amino acid substitutions per site.

As observed previously (9), the reported source of the genomes was not always consistent with the clade in which it was grouped. For instance, while publicly available genomes originally retrieved from human, baboon, elephant and cow gastrointestinal tracts were related to “*candidatus* Methanomethylophilus” (HA), this clade also contained MAGs derived from digestor and reactors (figure 1) reportedly not treating animal waste (table S1). Moreover, MAGs retrieved from pit mud of solid-state fermentation reactors used for the production of Chinese liquor were present in both the HA and FL clades (table S1). Similarly, “*ca.* M. intestinalis” Issoire-Mx1, *M.* luminyensis B10, and Methanomassiliicoccales archaeon RumEn M1, all retrieved from mammal hosts, grouped in the FL clade.

### Abundance of Methanomassiliicoccales clades differs in gastrointestinal and environmental samples

We assessed the abundance of species-level representative *Methanomassiliicoccales* taxa in publicly available metagenomes that included 145 samples from gastrointestinal tracts of non-human animals, such as cats, pigs, elks, cows, sheep, mice, white-throated woodrats, trouts, chickens and geese, and 160 environmental samples from sediment, ice, and diverse water and soil sources (table S2).

Taxa from all three clades were detected in a wide range of metagenomes from environmental and gut origin. We observed differences in environmental preference by clade. Abundance of taxa from Clade EX was highest on environmental metagenomes (0.001 % ± 0.0012) (figure 1). They were also detected in gut samples (0.0002 % ± 0.0005), albeit with a very low abundance in fecal (0.0003 % ± 0.0005), large intestine (0.0001 % ± 0.0002), stomach (0.0009 % ± 0.0006) metagenomes (figure 2). Given their low abundances, further analysis is focused on the FL and HA clades.

**Figure 2.**
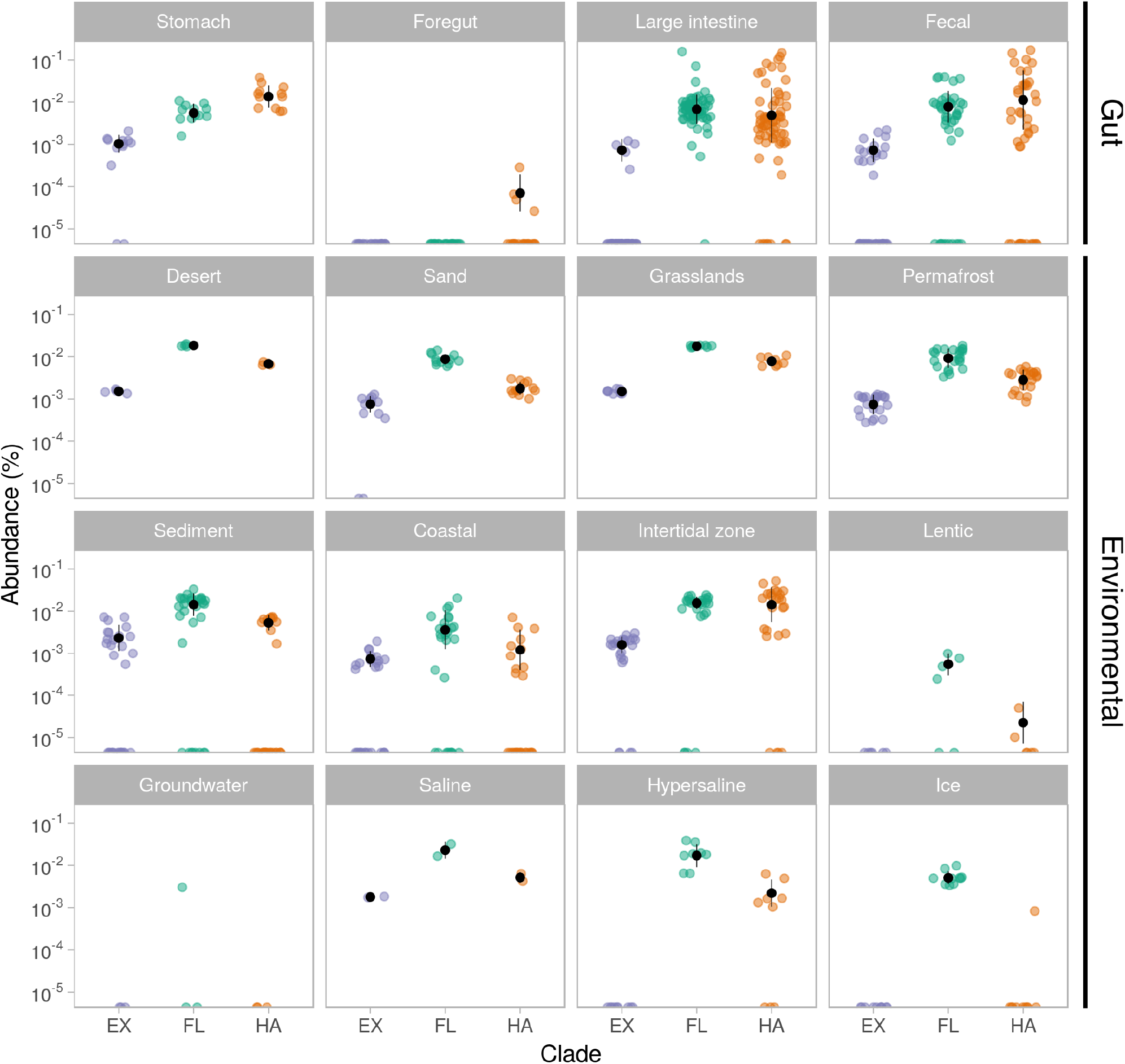
*Methanomassiliicoccales* clades are widespread but not abundant across a range of environments and animal hosts. Combined abundance of representative genomes of the EX (purple), FL (green), HA (orange) clades on metagenome samples from diverse biomes: stomach (n = 12), foregut (23), large intestine (66), fecal (44), desert (4), sand (12), grasslands (8), permafrost (22), sediment (31), coastal (28), intertidal zone (25), lentic (6), groundwater (3), saline (2), hypersaline (9) and Ice (10). Abundances calculated for individual genomes using KrakenUniq and aggregated by clade. Y-axis in logarithmic scale, black points indicate mean relative abundance in percentage, black bars indicate standard deviation.

The aggregated abundance of Clades FL and HA varied across biomes (figure 2). In agreement with their names, HA clade members were enriched in host-associated samples, and FL in non-host samples. The combined abundance of members of Clade FL was higher in samples from environmental biomes (0.01 % ± 0.008), although non-zero abundances were observed in digestive system metagenomes (0.008 % ± 0.015), with some samples containing levels comparable to that of Clade HA (figure 2).

The mean abundance of Clade HA in aggregate was higher in metagenomes from gut samples (0.014 % ± 0.03) compared to environmental biomes (0.004 % ± 0.008). However, among the environmental biomes, non-zero abundances of Clade HA were detected in freshwater (0.002 % ± 0.003), marine (0.006 % ± 0.011), saline and alkaline (0.002 % ± 0.002) and soil (0.004 % ± 0.003) samples.

We further validated the differences in clade abundances across biomes by generating a dendrogram of *Methanomassiliicoccales* taxa using the fold-change enrichment of individual taxa on gut versus environmental biomes, that is, the effect size of their enrichment in either direction. We then compared the structure of this dendrogram with that of the phylogenomic tree and found that they were positively correlated (cophenetic correlation = 0.67, P val. < 0.01).

Overall, we observed a low abundance of individual *Methanomassiliicoccales* taxa across all samples, ranging from 0 to 0.15 % (figure 2 and figure S1). The enrichment analysis of individual taxa from Clade FL on diverse biomes showed that while most were significantly enriched in environmental metagenomes, some taxa showed the opposite enrichment. *M. luminyensis* and Methanomassiliicoccus sp. UBA386 were not significantly enriched in gut or environmental biomes. “*ca.* M. intestinalis” Issoire-Mx1, Methanomassiliicoccales archaeon RumEn M1 and Methanomassiliicoccus sp. UBA6 were significantly enriched in gut biomes (figure 1), although they were also present in multiple environmental biomes (figure S1).

When assessed on a per-taxon basis, the vast majority of Clade HA taxa were significantly enriched in gut samples, with the exception of “*ca.* M. termitum”, which was highly abundant in soil samples from grasslands and water samples from intertidal zones (figure 1).

### Genome characteristics and core genes functions differ between Methanomassiliicoccales clades

Given the tendency of clades FL and HA to be enriched in environmental or animal metagenomes, respectively, we searched for genes and genome features linked to putative adaptations of *Methanomassiliicoccales* to an animal gut. For this, we compared 72 genomes from *Methanomassiliicoccales* taxa retrieved from humans, non-human animals and environmental sources.

We observed that genomes were more similar to others closely located on the phylogeny for genome GC content, genome length and total gene count (LIPA Adj. P < 0.01 in all cases) (figure 3). To determine whether these features differed between clades, while accounting for the autocorrelation due to evolutionary history, we performed a phylogenetic ANOVA. Clade FL taxa had significantly larger genomes (mean ± sd: 1985.1 Kb ± 245.1) than either the clades HA (1318.3 Kb ± 187.3) or EX (1872.2 Kb ± 173.8) (phylogenetic ANOVA Adj. P = 0.028). In accord, Clade FL also had the highest gene count (FL: 2153.1 genes ± 233.7; HA: 1377.7 genes ± 187.7; EX 1567.0 genes ± 90.5. Adj. P = 0.025). While non-significant, clades HA and EX taxa tended to have a lower GC content than Clade FL taxa (FL: 59.1 % ± 4.8; HA: 55.8 % ± 2.8; EX 54.4 % ± 0.5. Adj. P = 0.6).

**Figure 3.**
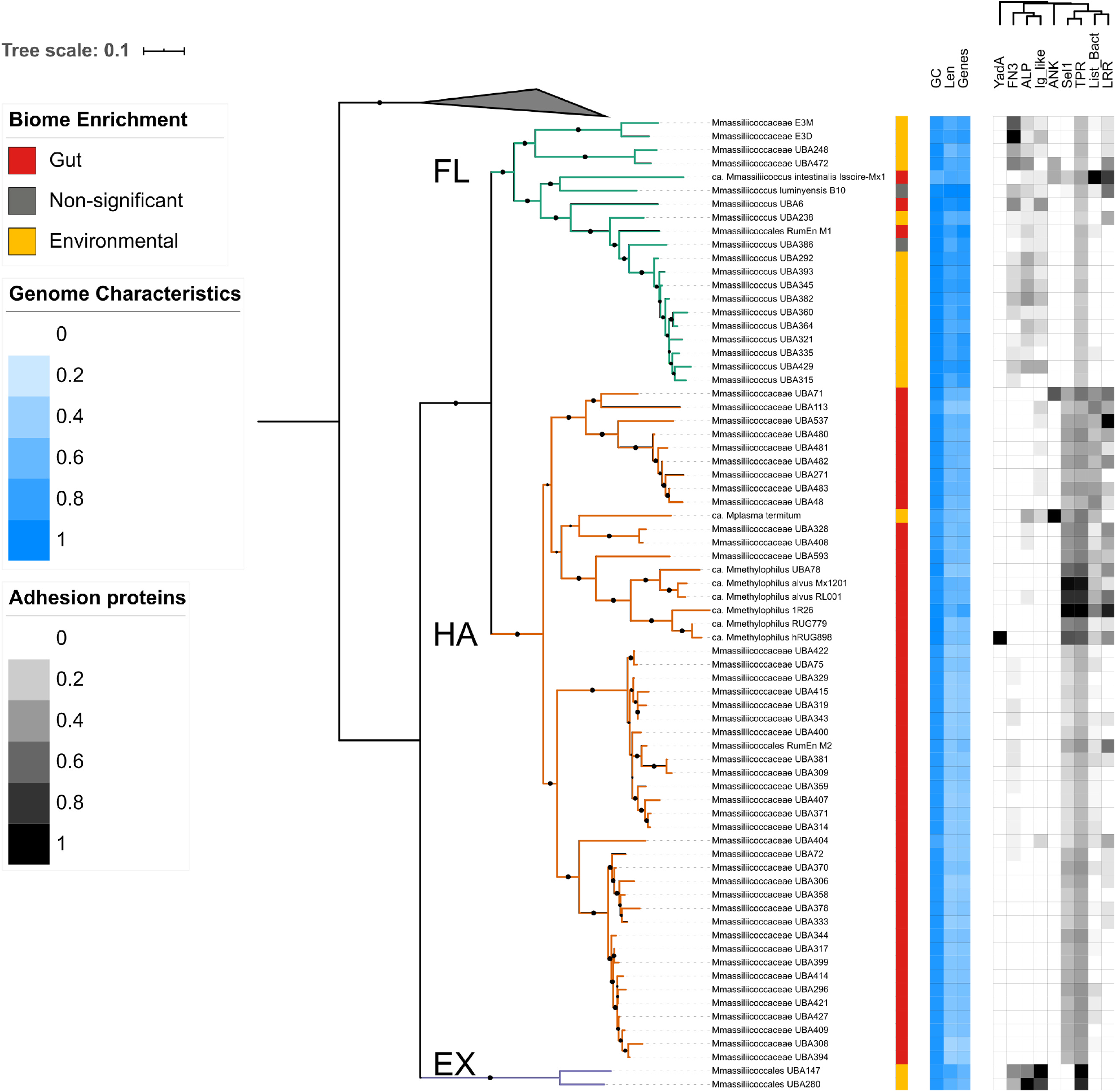
Genome characteristics and adhesion protein repertoire of *Methanomassiliicoccales* reflect division of the order into clades, although members of the Clade FL not enriched in environmental biomes resemble those of the Clade HA. The phylogeny is the same as shown in Figure 1. The colored strip summarizes the biome enrichment analysis. Heatmaps show genome features including genome GC content (GC; range: 41.26 %, 62.74 %), genome length (Len; 969.311 bp, 2.620.233 bp), and number of predicted genes (Genes; 1057, 2607) (blue scale); or repertoire of adhesion proteins: Sel1 containing proteins (Sel1; 0, 29), Listeria-Bacteroides repeat containing proteins (List-Bact; 0, 26), tetratricopeptide repeats (TPR; 7, 40), Ankyrin repeats (ANK; 0, 3), Leucine-rich repeats (LRR; 0, 9), Fibronectin type III (FN3; 0, 20) domains, Bacterial Ig-like domains (Ig-like; 0, 12), YadA-like domain (YadA; 0, 1) and adhesin-like proteins (ALP; 0, 12) (gray scale; columns ordered by hierarchical clustering).On both heatmaps the color intensity of each feature is relative to the maximum value of each category. Scale bar represent the number of amino acid substitutions per site.

To compare gene presence and absence across clades, we performed a pangenome analysis. After identification of orthologous gene clusters based on sequence similarity using PanX, we obtained 13,695 clusters, of which 7,312 were present at least once in Clade FL, 6,592 in Clade HA, and 1,833 in Clade EX. A large proportion of gene clusters were of unknown function according to the COG functional classification (38.4 % ± 4.3); gene clusters of unknown function tended to be small, with only one or two genes (figure S2 A, B). Principal component (PC) analysis of gene cluster presence/absence clearly differentiated clades along PC1.

We defined outlier taxa as FL taxa enriched in gut biomes (Methanomassiliicoccales archaeon RumEn M1, Methanomassiliicoccus sp. UBA6, “ca. Methanomassiliicoccus intestinalis” Issoire-Mx1, *Methanomassiliicoccus luminyensis* B10 and Methanomassiliicoccus sp. UBA386) and the HA taxon enriched in non-host biomes (“*ca.* M. termitum”). Outliers mostly clustered with their close relatives, not with the taxa enriched in the same biome (figure 4), with the exception of “ca. Methanomassiliicoccus intestinalis” Issoire-Mx1, which did not cluster with either clade.

**Figure 4.**
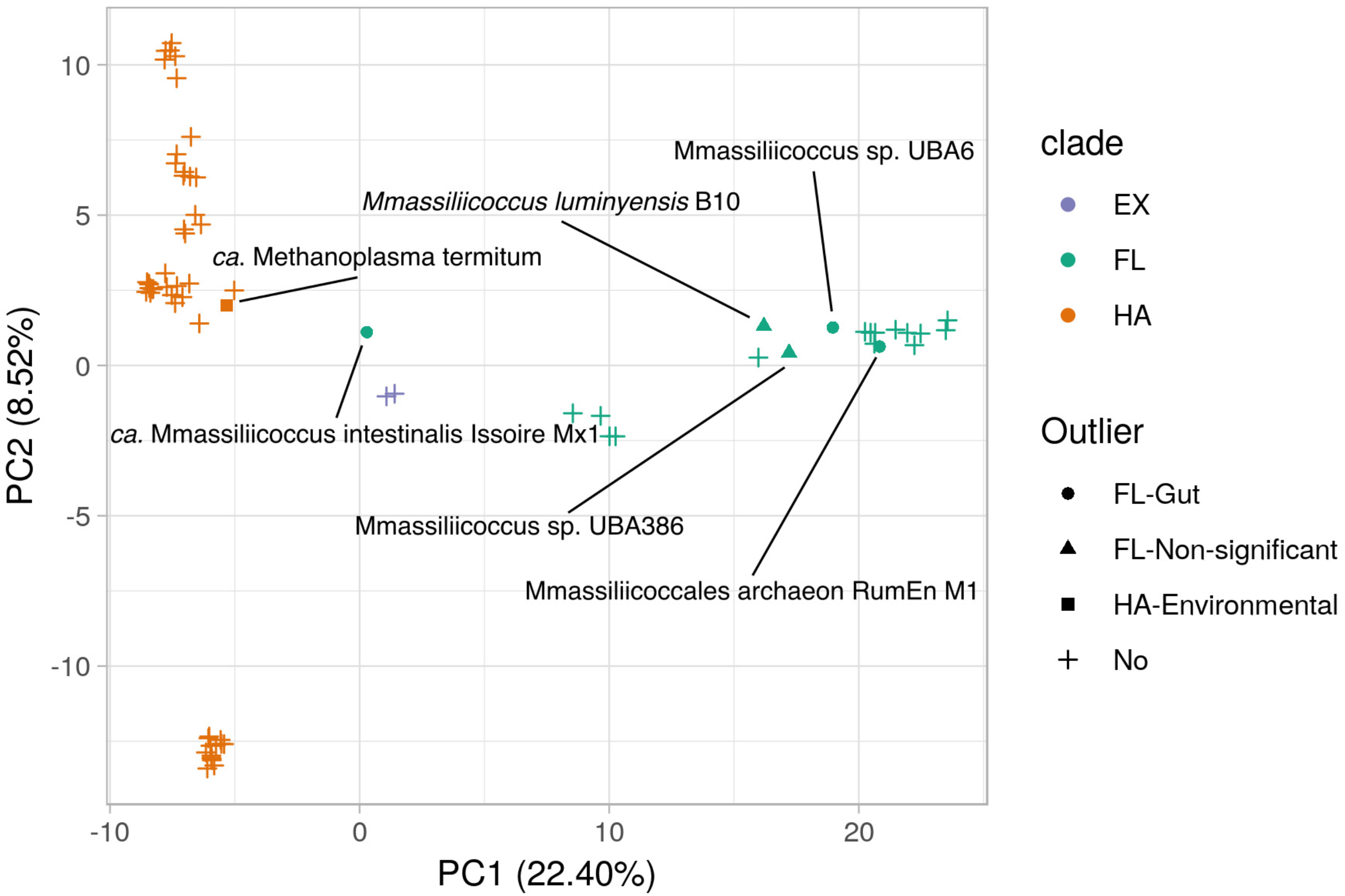
Ordination of gene content of *Methanomassiliicoccales* group taxa by phylogenetic clade rather than by biome enrichment. Principal component analysis of the gene cluster presence of taxa from clades FL (green), HA (orange) and EX (purple). Highlighted points correspond to outliers: taxa either not significantly enriched in environmental or gut biomes, or with enrichment opposite to the expectation given their clade.

### Gene clusters enriched in Clade HA evidence adaptation to the gut environment

Because of the small number of genomes that cluster within the Clade EX, and because they are largely absent from animal-associated samples, subsequent analyses focus on comparisons between the clades FL and HA.

To identify gene clusters potentially involved in the adaptation of members of Clade HA to a host environment, we compared the gene cluster content between clades. The gene cluster frequency spectrum shows many clusters with a small number of genes: 7,990 (58.3 %) gene clusters were singletons and 2,002 (14.6 %) were doubletons (figure S2 A, B). After removing rare gene clusters by filtering those with near zero variance, we included 2937 clusters, which we then used to perform in phylogenetic ANOVAs. Results reveal 14 gene clusters significantly enriched in

HA compared to FL (Adj. P < 0.1 in all cases). Three gene clusters are involved in detoxification and xenobiotic metabolism, namely, bile acid:sodium symporter (InterPro accession IPR002657), bleomycin resistance protein (IPR029068) and HAD-superfamily hydrolase (IPR006357). Two clusters are related to shikimate or chorismate metabolism: chorismate mutase II (IPR002701) and prephenate dehydratase (IPR001086). Other annotated clusters include the small unit of exonuclease VII (IPR003761), holliday junction resolvase Hjc (IPR002732), nitrogen regulatory protein PII (IPR015867), xylose isomerase-like protein (IPR013022) and metal-binding domain containing protein (IPR019271). Four had poor or no annotation (table 1).

**Table 1.** InterPro, eggNOG and Prokka annotations of gene clusters significantly enriched on clade HA compared to Clade FL. Four gene clusters with no annotation were omitted.

| Interpro accession | Interpro annotation | NOG accession | COG category | Prokka gene name | Prokka annotation |
| --- | --- | --- | --- | --- | --- |
| IPR001086 | Prephenate dehydratase | COG0077@NOG | E | pheA | Prephenate dehydratase |
| IPR002657 | Bile acid:sodium symporter/arsenical resistance protein Acr3 | COG0385@NOG | S | - | hypothetical protein |
| IPR002701 | Chorismate mutase II, prokaryotic-type | COG1605@NOG | E | aroQ | Chorismate mutase |
| IPR002732 | Holliday junction resolvase Hjc | COG1591@NOG | L | rutD | Putative aminoacrylate hydrolase RutD |
| IPR003761 | Exonuclease VII, small subunit | COG1722@NOG | L | xseB | Exodeoxyribonuclease 7 small subunit |
| IPR006357 | HAD-superfamily hydrolase, subfamily IIA | COG0647@NOG | G | gph | Glyceraldehyde 3-phosphate phosphatase |
| IPR013022 | Xylose isomerase-like, TIM barrel domain | 11IHC@NOG | L | - | hypothetical protein |
| IPR015867 | Nitrogen regulatory protein PII/ATP phosphoribosyltransferase, C-terminal | COG3323@NOG | S | - | hypothetical protein |
| IPR019271 | Protein of unknown function DUF2284, metal-binding | 11RTN@NOG | S | - | hypothetical protein |
| IPR029068 | Glyoxalase/Bleomycin resistance protein/Dihydroxybiphenyl dioxygenase | - | X | - | hypothetical protein |

### Genomic adaptations to the gut of members of the FL clade

To determine whether outlier taxa belonging to Clade FL had similar adaptations to the gut as members of Clade HA, we explored gene clusters present in these outliers and in Clade HA but that were rare in other members of Clade FL. We selected gene clusters present in the core genome of Clade HA (i.e. present in >80 % of taxa from this clade, see supplementary results) and present in less than half of FL taxa. A total of 15 gene clusters were obtained, most of them encoded by only one of the outlier taxa. Two gene clusters, ferrous iron transport proteins A and B (IPR030389 and IPR007167), were present in three of the outliers (Methanomassiliicoccus luminyensis B10, “*candidatus* Methanomassiliicoccus intestinalis” Issoire-Mx1 and Methanomassiliicoccales archaeon RumEn M1). Other clusters detected in more than one outlier included an uncharacterised membrane protein (IPR005182, in Methanomassiliicoccus sp. UBA6 and Methanomassiliicoccales archaeon RumEn M1), a putative nickel-responsive regulator (IPR014864, in Methanomassiliicoccus luminyensis B10 and Methanomassiliicoccus sp UBA386), and an ABC transporter (IPR037294, in Methanomassiliicoccus luminyensis B10 and Methanomassiliicoccus sp UBA386). The remaining gene clusters, detected once, corresponded to transcriptional regulators or proteins of unknown function.

### The genomes of taxa from clades HA and FL encode distinct repertoires of adhesion proteins

We compared between FL and HA clades two large groups of membrane proteins involved in adhesion: eukaryote-like proteins (ELPs), a series of protein families involved in microbial adherence to its host (38), and adhesin-like proteins (ALPs), a class of proteins hypothesized to be involved in the microbe-microbe interactions of *Methanobacteriales* in the gut (13). We aggregated the counts of gene clusters annotated as the ALP and ELP classes, and performed phylogenetic ANOVA. This analysis showed that members of each clade tended to encode a different repertoire of adhesion proteins (figure 3). Taxa from Clade HA had a higher mean count of tetratricopeptide repeats (Mean±SD count; HA: 16.30±06.56, FL: 9.55±1.70), Sel1 repeats (HA: 9.32±5.69, FL: 0.35±1.35), Listeria-Bacteroides repeats (HA: 3.68±3.76, FL: 1.65±5.78) and leucine-rich repeats (HA: 1.5±2.15, FL: 1.1±2.02) than FL taxa, although we did not observe significant differences in their frequency (Adj. P > 0.1 in all cases). Conversely, ALPs (FL: 2.25±1.48, HA: 0.14±0.61) and Ig-like domains, (FL: 1.55±1.32, HA: 0.20±0.53) tended to be more abundant in the genomes of members of Clade FL.

Interestingly, outlier taxa from Clade FL had gene counts of several of the adhesion factors higher than the mean of their own clade and more characteristic of clade HA. In some cases, the gene counts were higher than the mean for Clade HA. These included Listeria-Bacteroides repeats (gene cluster count - *M. luminyensis*: 2, “*ca.* M. intestinalis” Issoire-Mx1: 26, Methanomassiliicoccales archaeon RumEn M1: 2), Sel1 repeats (*M. luminyensis:* 1, “*ca.* M. intestinalis” Issoire-Mx1: 6), and leucine-rich repeats (*M. luminyensis:* 5, “*ca.* M. intestinalis” Issoire-Mx1: 7).

### Methanomassiliicoccales taxa cooccur with each other, with other Archaea, and with TMA producing bacteria in the human gut

We characterized the distribution of *Methanomassiliicoccales* across a collection of human gut metagenomes derived from 34 studies. Together, the combined 4472 samples represented people from 22 countries, resulting in 35 unique datasets (*i.e.*, study-country combination). Across the whole set, we detected just two genera: *Methanomassiliicoccus* (Clade FL) and “*ca.* Methanomethylophilus” (Clade HA), both rare members of the human gut microbiota (figure 5). “*ca.* Methanomethylophilus” was detectable in 19 out of 35 datasets; on these 19 datasets it had a prevalence ranging from 0.5 % to 41.7 %, and mean abundance ranged from 4.8*10^−6^ % to 2.2*10^−2^ %. Similarly, *Methanomassiliicoccus* was detectable in 22 of the 35 datasets; on the 22 datasets it had a prevalence range of 1 % to 25.7 % and a mean abundance range of 1.5*10^−5^ % to 1.0*10^−2^ % (table S4).

**Figure 5.**
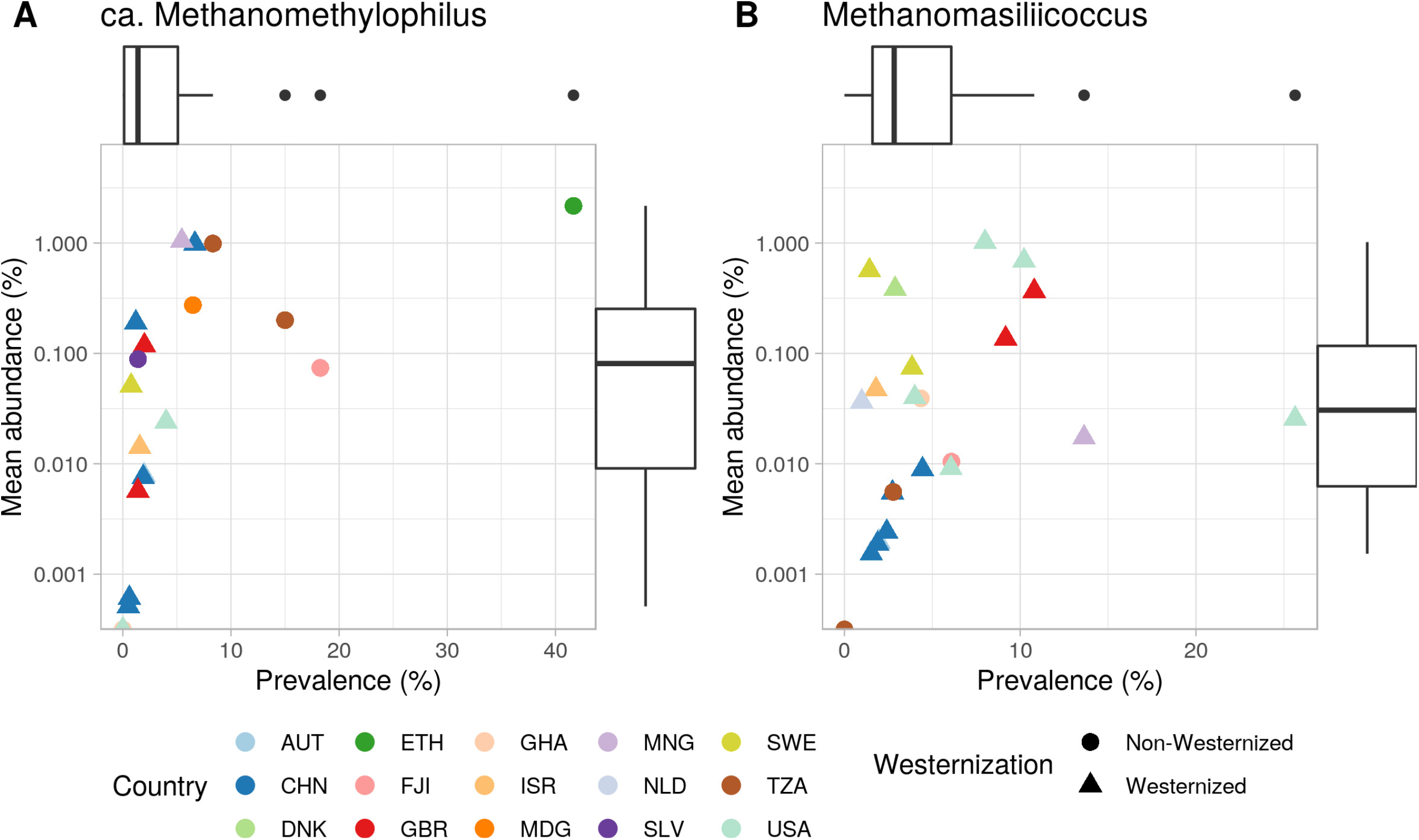
Members of *Methanomassiliicoccales* are rare members of the human gut microbiota. Scatter plots of the genera A) *ca.* Methanomethylophilus and B) *Methanomassiliicoccus* show that their prevalence and mean abundance is low across most studies and populations (n = 4472; 35 datasets) with subjects from Australia (AUT), China (CHN), Denmark (DNK), Ethiopia (ETH), Fijo (FJI), Great Britain (GBR), Ghana (GHA), Israel (ISR), Madagascar (MDG), Mongolia (MNG), The Netherlands (NDL), El Salvador (SLV), Sweden (SWE), Tanzania (TZA) and the United States (USA).

We tested associations of these two genera with age, sex and westernization status of the subjects using linear mixed models that included the dataset and country as random effects. Subjects from non-westernized countries had a significantly higher prevalence of “*ca.* Methanomethylophilus” (mean prevalence ± SD: Non-westernized = 8.9 % ± 28.5, Westernized 1.1 % ± 10.3; P Adj. = 0.002). Westernized individuals were more likely to harbor higher *Methanomassiliicoccus*, although differences were not significant (Non-westernized = 3.9 % ± 19.4, Westernized 5.0 % ± 21.7; P Adj. > 0.1). The age and sex of the individuals did not explain variance in the prevalence or abundance of either genus (Adj. P > 0.1 in all cases).

To identify other microbial taxa positively associated with members of *Methanomassiliicoccales* in the human gut, we calculated a network of positively associated microorganisms (*i.e.* coabundant taxa) across samples (rho >0.1 in all cases) (53). In addition, we determined which taxa were present with members of *Methanomassiliicoccales* more than expected by chance (*i.e.* cooccurring taxa) relative to a permuted null model (43). Results showed that both “*ca.* Methanomethylophilus” and *Methanomassiliicoccus* were part of the same coabundance network, together with a third archaeal genus, *Methanoculleus (*order *Methanomicrobiales*). We did not find evidence of positive or negative abundance associations of either *Methanomassiliicoccales* genus with *Methanobrevibacter.* Coocurrence analysis showed a random association pattern between these taxa (P val. > 0.05 for both “*ca.* Methanomethylophilus” and *Methanomassiliicoccus)*, indicating that their ecological niches do not overlap with *Methanobrevibacter.*

Analysis of the combined network of “*ca.* Methanomethylophilus” and *Methanomassiliicoccus* revealed a large overlap between taxa associated with either genus (figure 6): out of 119 taxa in the network, 86 (72.3 %) were associated with both. Moreover, 51 taxa (42.9 %) also had a significant positive cooccurrence pattern with both genera (P val. < 0.05 in all cases). Most bacterial members of this network had an overall low relative abundance. Interestingly, they included several taxa whose genomes contain genes encoding enzymes involved in TMA production, including *Bacteroides, Campylobacter, Yokenella, Mobiluncus, Proteus, Providencia* and *Edwardsiella* (54).

**Figure 6.**
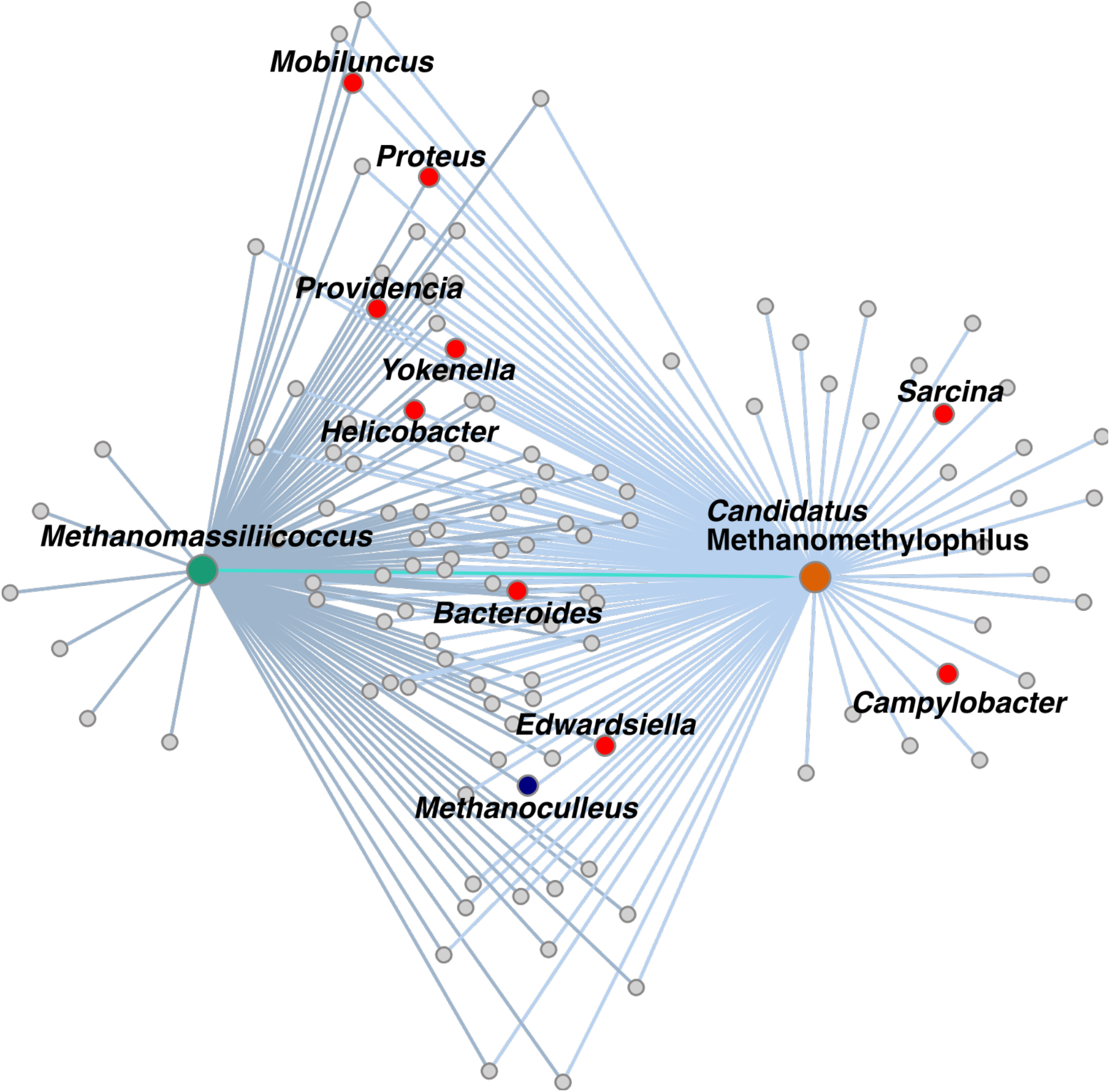
Coabundance networks of *Methanomassiliicoccus* (green node, dark edges) and “*ca.* Methanomethylophilus” (orange node, light edges) in the human gut largely overlap. Both *Methanomassiliicoccales* genera are significantly co-abundant (cyan edge). Their abundances are also coordinated with another archaeon (blue node) and TMA-producing bacterial taxa (red nodes).

### Abundance of Methanomassiliicoccales is not concordant in monozygotic or dizygotic human twins

To evaluate whether host genetics influences the abundance of *Methanomassiliicoccales* in the human gut, we compared the intraclass correlation coefficient (ICC) of their abundances at the genus level using a set of 153 monozygotic (MZ) and 200 dizygotic (DZ) twin pairs from the TwinsUK cohort. As control, we first compared the mean ICC across all taxa between MZ and DZ twins, and found that ICC_MZ_ (0.1) was significantly higher than ICC_DZ_ (0.03) (P val. < 0.01). In addition, we assessed the ICC values of bacterial (*Christensenella, Faecalibacterium and Bifidobacterium*) and archaeal (*Methanobrevibacter*), and consistently found a higher correlation on MZ compared to DZ twins (table S5). We were only able to assess ICC values of *Methanomassiliicoccus*, as it was the only *Methanomassiliicoccales* detected in the twins with a prevalence (8.64 %) above the 5 % cutoff (see methods). We did not detect a significant concordance between the abundances of *Methanomassiliicoccus* in MZ (ICC_MZ_ = 0.004, Adj. P = 0.59) or in DZ twins (ICC_DZ_ = 0.017, Adj. P = 0.71). Given the low abundance of *Methanomassiliicoccales* taxa, we performed a sensitivity analysis using samples with a high sequencing depth (>12 million reads/sample), however, we did not observe differences in the abundance and prevalence of the *Methanomassiliicoccales* genera nor the ICC estimates (not shown).

## Discussion

While the source of the members of the *Methanomassiliicoccales* has been noted in previous surveys of single markers such as 16S rRNA and mcrA genes (9, 10), here we searched metagenomes from host associated and environmental samples for their relative abundances. Overall, the HA taxa were enriched in host associated samples and the FL taxa in environmental samples; intriguingly, all taxa regardless of clade were detected in both biomes. This suggests that members of the order *Methanomassiliicoccales* are generalists with an overall habitat preference according to clade, although there were some exceptions to the general pattern. We show that members of *Methanomassiliicoccales* use many of the same adaptations to the gut as other methanogens. These adaptations include genome reduction, and genes involved in the shikimate pathway and bile resistance. In addition, gut-enriched taxa possess a distinct repertoire of genes encoding adhesion factors. We observed that potential adaptations to the gut differed by clade, not preferred habitat, indicating convergence on a shared niche through different genomic solutions. In the human gut, *Methanomassiliicoccales* taxa correlated with TMA-producing bacteria, rather than host genetics or other host factors.

For members of the HA clade, adaptations to life in the gut included an enrichment of genes involved in bile acid transport, efflux pumps, and hydrolases, which play a role in tolerance to these compounds in the gastrointestinal tract (55). This adaptation is also shared with other members of the gut microbiota, including *Methanobacteriales: M. smithii* and *M. stadtmanae* are resistant to bile salts (2, 3). Other gene clusters with known function enriched in Clade HA are involved in metabolism of shikimate and chorismate. The shikimate pathway is involved in the synthesis of aromatic amino acids in plants and microbes, but is absent in mammals. Shikimate metabolism is carried out by archaeal (56) and bacterial (57, 58) members of the animal gut microbiota, and was reported as one of the most conserved metabolic modules in a large-scale gene catalogue from the human gut (59). Derivatives from the aromatic amino acids are known to be bioactive in the mammal host (60).

In addition, members of the HA clade had a particular set of adhesion factors, known to be involved in the maintenance of syntrophic relationships of the methanogens with bacterial (12, 61) or eukaryotic (62) microorganisms. Two groups of adhesion factors, proteins containing Sel1 domains and Listeria-Bacteroides repeats, have been previously studied on *Methanomassiliicoccales* taxa retrieved from the gut (11, 52). Our assessment of these factors in the broader context of the order *Methanomassiliicoccales* showed that these two groups of proteins are characteristic of Clade HA rather than FL, with the exception of the outlier taxa. Indeed, while the repertoire of ELPs and ALPs differs between HA and FL taxa, it was similar between species inhabiting the gut regardless of their clade.

In contrast, members of Clade FL appear to be generalists that colonized the animal gut independently from the HA clade. It has been previously noted that *M. luminyensis*, an outlier from Clade FL, could have a facultative association to the animal gut. It possesses genes involved in nitrogen fixation, oxidative stress (11) and mercury methylation (52), which are common in soil microorganisms but rare in members of the gut microbiota (63). In accord, we observed that members of Clade FL are widespread and abundant on soil, water and gut metagenomes, with a preference for environmental biomes. Similarities in ELP content between gut-dwelling taxa from both clades indicate that interaction with the host or other members of the gut microbiota might be a key factor in the adaptation of these methanogens.

Analysis of the gene content of outlier taxa from Clade FL showed that they tended to be more similar to members of their own clade than to taxa from Clade HA, with the exception of “*ca*. M. intestinalis” Issoire-Mx1, which was distinct from either Clade FL and HA. In addition there was little overlap in gene clusters commonly observed in Clade HA and outlier taxa from Clade FL, with the exception of the adhesion factors discussed below. These observations support the hypothesis that colonization of animal guts by members of *Methanomassiliicoccales* occurred in two independent events (11, 52), and suggests that there is not one solution to life in the gut for these *Archaea*, as members from two clades seem to have solved the problem with a different set of adaptations.

Characterization of the abundance of *Methanomassiliicoccales* across human populations showed members of this group are rare in the microbiota of healthy adults. We did not detect them in all the studied populations, and when detected, they had low prevalence and abundance. Nevertheless, this analysis allowed us to assess whether *Archaea* in the human gut are mutually exclusive. We observed positive correlations of “*ca.* Methanomethylophilus” and *Methanomassiliicoccus* with each other and with *Methanoculleus*, another rare archaeal member of the gut microbiota (64). We did not find evidence of association between members of *Methanomassiliicoccales* and *Methanobrevibacter*, positive or otherwise, confirming the previous report that these methanogens are not mutually exclusive (46); abundance of H_2_ in the gut, together with differences in other substrate utilization, might result in non-overlapping niches (65).

While genus *Methanobrevibacter* has been consistently found to have a moderate heritability the TwinsUK (19, 46, 66) and other cohorts (48, 67), it was not the case for members of *Methanomassiliicoccales.* Similar to humans, methane production (68) and abundance of *Methanobrevibacter* (69) are also heritable in bovine cattle, but not *Methanomassiliicoccales* taxa (69). Thus, host genetics might be linked to particular taxa and methanogenesis pathways, not to all *Archaea* or methane production as a whole.

Genera “*ca.* Methanomethylophilus” and *Methanomassiliicoccus* cooccur with TMA-producing bacteria (54), further supporting their potential use as a way of targeting intestinal TMA (70). The exact nature of the ecological relationships each of these taxa establishes with other members of the microbiome remains to be elucidated. In a facilitation scenario between the methanogens and H_2_- and TMA-producers, freely available TMA and H_2_ required for methylotrophic methanogenesis could be utilized by *Methanomassiliicoccales* taxa (71), without cost to the producer. Alternatively, the methanogens could establish syntrophic interactions with other microorganisms, whereby the consumption of these metabolites is also beneficial to the producer (71).

The present study extends our understanding of the order *Methanomassiliicoccales* by revealing genomic adaptations to life in the gut by members of both clades that make up this group. Furthermore, the positive correlation between the relative abundances of these TMA-utilizing archaea with TMA-producing bacteria in the gut is a first step towards understanding how they may be harnessed for therapeutic management of gut TMA levels in the context of cardiovascular disease.

## Acknowledgements

This work was supported by the Max Planck Society. We thank EMBO, the organizers and participants of the Bioinformatics and genome analyses course held at the Fondazione Edmund Mach in San Michele all’Adige, Italy, for sponsoring the attendance of J.dlC-Z and for their feedback. We are also grateful to Daphne Welter, Jessica Sutter and Albane Ruaud for the fruitful discussions and comments. The study also received support from the National Institute for Health Research (NIHR) BioResource Clinical Research Facility and Biomedical Research Centre based at Guy’s and St Thomas’ NHS Foundation Trust and King’s College London. We declare no competing interests.

## Supplementary figure legends

**Figure S1. *Methanomassiliicoccales* taxa from all clades are widespread but not abundant across a range of environments and animal hosts.** The abundance of members of the FL and HA clades is comparable within similar biomes, in particular, animal derived metagenomes. Abundance of each representative genome on diverse metagenome and environmental metagenome samples colored by clade (FL: green, HA: orange, EX: purple). Abundances calculated for individual genomes using KrakenUniq and aggregated by clade. Note that the Y-axis is in logarithmic scale and each plot has a different scale. Black points indicate mean relative abundance in percentage, black bars indicate standard deviation. Metagenome samples from stomach (n=12), foregut (23), large intestine (66), fecal (44), desert (4), sand (12), grasslands (8), permafrost (22), sediment (31), coastal (28), intertidal zone (25), lentic (6), groundwater (3), saline (2), hypersaline (9) and Ice (10).

**Figure S2. Small clusters of unknown function dominate the pangenome of the order *Methanomassiliicoccales.*** Gene cluster frequency spectrum of the order Methanomassiliicoccales separated by (A) unknown or (B) known function. (C) Fraction of gene clusters belonging to each COG category per clade. Core clusters were defined as present in ≥ 80% of genomes of a clade; for the complete order, gene clusters were present in ≥ 80% of the included genomes and at least one member of each clade. The proportion of clusters of unknown functions in the core genome of each clade was large and varied between clades, ranging from 23.0 % in Clade HA to 38.5 % in Clade EX. The proportion of unknown clusters was lowest in the complete taxonomic order, where it only accounted for 14.7 % of gene clusters. COG functional classification descriptions by groups. Information Storage and processing: (B) Chromatin structure and dynamic, (J) Translation, ribosomal structure and biogenesis, (K) Transcription, (L) Replication, recombination and repair. Cellular processes and signaling: (D) Cell cycle control, cell division, chromosome partitioning, (M) Cell wall/membrane/envelope biogenesis, (N) Cell motility, (O) Post translational modification, protein turnover, chaperone, (T) Signal transduction mechanisms, (U) Intracellular trafficking, secretion, and vesicular, transport, (V) Defense mechanisms, (Z) Cytoskeleton. Metabolism: (C) Energy production and conversion, (E) Amino acid transport and metabolism, (F) Nucleotide transport and metabolism, (G) Carbohydrate transport and metabolism, (H) Coenzyme transport and metabolism, (I) Lipid transport and metabolism, (P) Inorganic ion transport and metabolism, (Q) Secondary metabolites biosynthesis, transport and catabolism. Poorly characterized: (X) No annotation retrieved, (S) Function unknown.

## Supplementary table legends

**Table S1.** CBI assembly accession, genome characteristics, study information and source of isolation of 71 publicly available genomes from the order Methanomassiliicoccales retrieved from NCBI in June 2018, plus the ca. M. alvus MAG here reported. Study accession and title of UBA genomes obtained from supplementary tables of Parks et al., 2017 (doi: 10.1038/s41564-017-0012-7), otherwise, obtained from NCBI bioproject.

**Table S2.** SRA and MGnify accession information of publicly available metagenome samples from gastrointestinal and environmental biomes

**Table S3.** SRA, study and country information of publicly available human gut metagenome samples

**Table S4.** Prevalence and mean abundance of candidatus Methanomethylophilus and Methanomassiliicoccus across multiple human populations.

**Table S5.** Intraclass correlation coefficients (ICC) of and FRD-adjusted P values of relative abundances Methanomassiliicoccus and other control taxa on monozygotic and dizygotic twins

